# Neurocognitive Mechanism of Radiologists’ Perceptual Errors: Results of Preliminary Studies

**DOI:** 10.1101/2024.04.29.591746

**Authors:** Michael A. Bruno, Elizabeth A. Krupinski, Scott C. Bunce, Grayson L. Baird, Caitlin Mills, Prasanna R. Karunanayaka, Howard Egeth, Ryan Chang, Ross Cottrill, Samuel Jump, Krishnankutty Sathian, Timothy J. Mosher

**Affiliations:** Department of Radiology, Penn State Health Milton S. Hershey Medical Center, Hershey, PA; Penn State College of Medicine, Hershey, PA; Department of Radiology, Emory University School of Medicine, Atlanta, GA; Department of Radiology, Rhode Island Hospital, Brown University, Providence, RI; Department of Educational Psychology, University of Minnesota Twin Cities, Minneapolis, MN; Department of Psychological & Brain Sciences, Johns Hopkins University, Baltimore, MD; Department of Neurology, Penn State Health Milton S. Hershey Medical Center, Hershey PA

**Keywords:** Functional Connectivity, Network Neuroscience, Radiologist Performance, Perceptual Error

## Abstract

**Background:** The most prevalent type of radiologist error is failing to detect abnormalities on images, the so-called “perceptual error.” The prevalence of this type of false-negative (FN) error remains essentially unchanged since it was first described in 1949.

**Purpose:** The purpose of this research is to identify a potential neurocognitive mechanism contributing to radiologists’ susceptibility to perceptual error, in order to inform intervention strategies to reduce such errors in practice. These experiments evaluated the relationship between brain network activation states and radiologists’ perceptual errors on two distinct visual tasks utilizing functional MRI (*f*MRI) and functional Near Infrared Spectroscopy (*f*NIRs).

**Materials and Methods:** A prospective study consisting of three experiments was carried out on a small number of radiologist subjects. The first two experiments used *f*MRI, with participants performing two distinct types of visual tasks, respectively: the first was a task requiring subjects’ continuous attention and the second task required visual search. For the first of these experiments, simultaneous functional Near-Infrared Spectroscopic Imaging (*f*NIRs) was utilized along with *f*MRI. The second experiment was combined *f*MRI and eye-tracking. A third experiment using *f*NIRs alone was an observational study of subjects’ neurocognitive states during their usual practice.

**Results:** An approximately threefold increased risk of FN perceptual errors (misses) was observed in the presence of a particular error-prone neurocognitive state (EPS) involving simultaneous co-activation of elements of the Default Mode Network (DMN) and Frontoparietal Network (FPN), which was detectable by both functional imaging modalities, with high concordance. EPS episodes appeared to be stochastic in occurrence, and occurred without operator awareness. We also found a high prevalence of the EPS in radiologists performing their normal interpretive tasks in their actual practice setting.

**Conclusion:** Our results suggest that dynamic interactions between brain networks leading to a particular error-prone state (EPS) may underlie a substantial fraction of radiologists’ perceptual errors. We demonstrate that this EPS can be detected unobtrusively in the clinical setting. These results suggest potential intervention strategies for perceptual error, the largest class of radiologist errors in practice.

**Key Results/Highlights:**

1. Periodic episodes of a discrete neurocognitive state were observed in radiologists during specific visual tasks and in actual clinical settings.
2. There was an approximately threefold increased risk of perceptual error during this state. Most FN errors for the two visual tasks occurred during these brief episodes (p < 0.01).
3. There was also a highly significant anti-correlation of the prevalence of the error-prone neurocognitive state (EPS) with subject age (p < 0.001).

**Conflicts:** The authors report no conflicts of interest or potential competing interests.

**Summary Statement:** We report experimental results corelating perceptual errors by radiologists to episodic fluctuations in brain network activation, which appear to occur on a stochastic basis. These produce an error-prone neurocognitive state outside of operator awareness or control that is associated with an approximately threefold increase in the risk of perceptual error.

## Introduction

Radiologists frequently fail to visually perceive abnormalities that are later shown to have been present on images, including many that are readily detected in retrospect. This type of error, known as “perceptual error,” is by far the most common error type for radiologists, accounting for the majority of radiologists’ errors in practice (1, 2). In contrast, relatively few diagnostic errors are due to radiologists’ misinterpretation of detected findings, cognitive biases or clinical knowledge deficits (3, 4). Since first described by Garland in the late 1940’s (5), the prevalence of perceptual error in radiology has not appreciably changed (6). While radiologist error rates as low as 3-4% have been reported in the setting of low prevalence of positive findings, higher error rates, on the order of 30% are typical when radiologists are presented with test cases that have a higher prevalence of abnormalities, with some studies showing up to 70% of lung cancers missed on chest x-rays and 39% on CT scans (7). The etiology of these common errors of omission is likely the result of both visuo-perceptual and cognitive factors acting together in complex ways (8). Such errors appear to be due to a basic human factor, as they have not been mitigated by substantial improvements in imaging technology and radiological knowledge over the past several decades.

It is an attractive hypothesis that many radiologists’ perceptual errors could be the result of natural fluctuations in attention or task engagement related to human operator neurocognitive states that lead to momentary lapses in attention or receptiveness to visual stimuli. Critically, such inattentive states can occur without operator awareness (9), making them difficult or impossible to detect in real-time, although they are often readily apparent in retrospect (see figure 1). When such errors involve highly conspicuous abnormalities, radiologists are often at a loss to explain how such a finding could have been missed. When errors of this type lead to protracted delays in diagnosis, they can have devastating consequences for patients. For example, in patients with many types of cancer, delays in diagnosis can greatly restrict options for treatment and lead to worsened outcomes.

**Figure 1.**
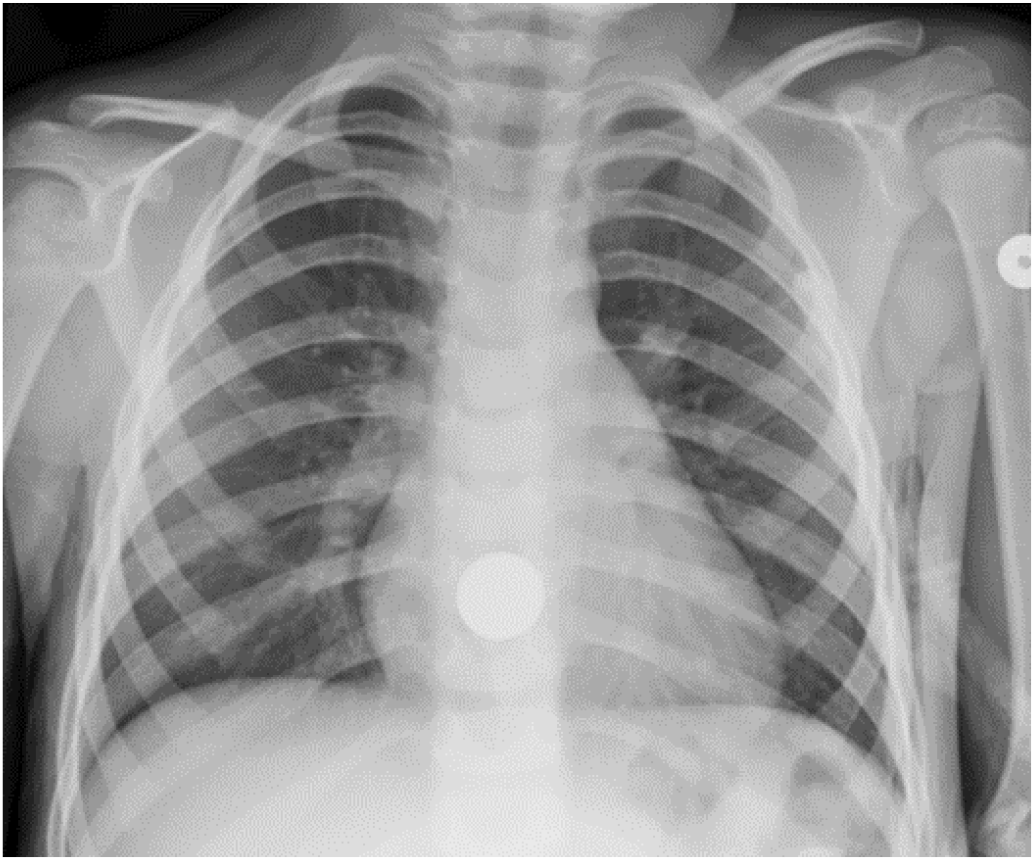
A chest radiograph of a child obtained to rule out a swallowed coin. The presence of the coin within the esophagus is readily apparent on the image, but was missed twice by a highly experienced and skilled pediatric radiologist. This case illustrates that even very conspicuous abnormalities can be overlooked by highly skilled observers (from Reference 2, used with permission).

An emerging literature within the broader field of network neuroscience has described a particular neurocognitive state that results from interactions between two well-described networks of cortical brain regions (10, 11). The first is the “Default Mode Network,” or DMN, which was first identified by Raichle *et al*. (12, 13), and which is believed to underly internally-focused modes of thought, and the other is the Frontro-parietal network (FPN), which is believed to underly externally-directed thought and tasks requiring attention to outside stimuli (14).

It has been observed that the DMN generally acts in opposition to the FPN, with the two networks alternating in periods of activation and suppression during the course of a normal day. Extensive observational research using BOLD *f*MRI has shown that, under usual circumstances, activity within either one of these two networks is anti-correlated with activity in the other. Indeed, human subjects are observed to “toggle” between them as they begin and finish externally-directed tasks. The “toggle” function is apparently under control of additional cortical networks (15). Researchers have, however, identified a particular episodic or transient neurocognitive state wherein there is simultaneous co-activation of these two functional brain networks, creating a hybrid DMN+FPN neurocognitive state. It has further been shown that this state is associated with an apparent decoupling of sensory inputs from cognitive awareness (16). This state can occupy up to 30% of a person’s waking hours and is generally not under volitional control (17, 18, 19, 20). Subjects are notably less receptive to visual stimuli at these times, despite the appearance of continuing to attend to and search the input (11, 21, 22, 23).

This research aimed to test the hypothesis that transient episodes of this well-described hybrid neurocognitive state increase the risk of perceptual error in radiologists performing visual tasks, possibly due to its associated phenomenon of perceptual decoupling. In this work we refer to this neurocognitive phenomenon as the “Error-Prone State,” or EPS, and suggest that it might provide a mechanistic explanation for many (perhaps a majority) of radiologists’ perceptual errors in practice.

To test this hypothesis, we have designed three experiments that will evaluate the occurrence of the EPS in experienced radiologists in two very distinct visual tasks, one involving continuous subject attention and another involving visual search, as well as in the radiologists’ actual work setting, with the goal of determining whether EPS is a significant risk factor for perceptual error in our subjects, to determine the prevalence of EPS under each of these conditions, and to evaluate the feasibility of detecting EPS using a relatively unobtrusive measure as a path for future work aimed at interventions to reduce perceptual errors.

## Materials and Methods

Our protocol was IRB approved. Three experiments were conducted in our pilot study, utilizing functional MRI (*f*MRI) and functional near-infrared spectroscopic imaging (*f*NIRs) to assess brain states of our subjects as they carried out various cognitive tasks. For the experiments involving *f*NIRS we received a free loan of *f*NIRs equipment along with customized analytical software from the NIRx Corporation (Berlin, Germany). The authors retained full control of the data and determined all of the content which has been submitted for publication.

Three experiments were carried out. In the first experiment, 10 radiologists performed a visual task requiring their continuous attention for 15 minutes, divided into two 450-second sessions. Subjects included 8 males and 2 females, ranging in age from 37-65 (mean = 49.5). All but one were right-handed. The subjects were instructed to watch a small, gray disc as it slowly meandered across their viewing screen on a textured grayscale background reminiscent of the appearance of x-ray images. At pseudo-random times, and without warning, the luminosity of the disc abruptly changed, at which time they were to press a button indicating that they had detected an event. By design, this continuous attention task (CAT) uses subtle luminosity changes. These visual events are readily detectable, but only if the subject is continuously tracking the disk and paying attention to it. The task is designed such that it is virtually impossible for a subject to determine whether a change-event occurred retrospectively. The single visual target is always in plain view and moves across the screen very slowly, such that the task does not involve visual search. When the subject detects a change, they signal by pushing a button, allowing measurement of response times. By measuring accuracy, *e*.*g*., successful detection and response times, it is possible to detect lapses in the subjects’ attention with high temporal precision, and correct responses were detected in as little as 550 ms. As a cut-off value, any response recorded within 3 seconds of an event was accepted; after that time, any further-delayed response was counted as a miss. Responses that were not temporally related to an event were counted as FP. The task was not in itself difficult or challenging, but required the continuous attention of the subject in order to detect the visual events. It would be essentially impossible for a subject to be certain that a change had occurred after the fact, *i*.*e*., if their attention to the disk was interrupted or intermittent.

For assessment of subjects’ neurocognitive states during the task, we utilized simultaneous whole-brain *f*MRI and *f*NIRs of the prefrontal cortex, using an MRI-compatible *f*NIRs system (NIRx Scout, NirX Medical Technologies, LLC, Berlin). *f*MRI was performed using a 3.0T Siemens PrismaFit MRI system (Siemens, Erlangen, Germany). Technical details of our experimental and analytical approach, as well as a discussion of the *f*NIRs technology that was used, are included in the appended technical supplement. For the *f*MRI analysis, the time course of activation BOLD (OxyHb) signal was corelated for the DMN and FPN to compare temporal behaviors to detect co-activation events, defined as a positive (in-phase) dynamic correlation between the two normally out-of-phase (anticorrelated) networks using scale-independent amplitudes of activity. Independent analysis was performed for the 8 of these subjects for which *f*NIRs data were also available, and *f*NIRs and *f*MRI results were compared for these 8 subjects in order to validate the use of *f*NIRs, which was limited to the PFC, against the whole-brain *f*MRI “gold standard.”

Each 450-second run had 50 visual events to be detected, with a pseudorandom variable inter-stimulus interval. The timing of FN “miss” errors was correlated to both the *f*MRI and *f*NIRs to ascertain to what extent subject errors correlated with the “target” EPS, and to assess the level of concordance between the two techniques, with the goal to validate the later use of only *f*NIRs in the clinical setting. The rationale for using *f*NIRs, which is limited to visualizing BOLD signal only the relatively superficial PFC components of the DMN and FPN was based on the well-established observation that the several brain areas that comprise each network demonstrate tightly-coupled, coordinated patterns of activation, such that it would be unusual for some components of either the DMN or FPN to be active when others are not; this is a fundamental feature defining these networks (12, 13). We therefore predict that it would be most likely that activation of the PFC components of the DMN and FPN alone would be highly predictive of the behavior of the entire network. The degree of correlation was measurable by comparison of the simultaneously-acquired and independently analyzed *f*NIRs and *f*MRI data.

The second experiment utilized *f*MRI alone (no *f*NIRs) with the addition of concurrent eye-tracking, using a conventional psychophysical visual search task. This experiment was designed to determine whether the association of EPS and error differed when the subjects were actively searching the images for a target, in order to generalize the results beyond sustained attention. Visual search is fundamental to the practice of radiology. Eye-tracking was performed using an SR Eyelink™ system (SR Research, Ottaway, ON, Canada) in order to determine whether or not the subjects looked at the targets with their central, foveal vision as part of their decision-making process. A total of 6 Radiologist subjects (five male, one female, all right-handed) were tasked with detection of a target grayscale “T” on a textured grayscale background among numerous “L” distractors (Figure 2). In preparatory work using multiple radiologist and non-radiologist volunteers, the task was designed to be moderately difficult (on the basis of both objective subject performance and subjective measures). Task difficulty was adjusted by manipulation of the opacity/translucency of the T’s and L’s, altering the number of “L” distractors, as well as limiting the time subjects were allowed to view each task to 10 seconds. Subjects were engaged for approximately 15 minutes in two 450-second sessions. Each trial contained multiple “L” distractors and may or may not have included a “T” target. Each 450-second session included 60 T/L observational frames. If a “T” was detected, the subject would indicate this by pressing a button, and they could also indicate when they believed that no “T” was present, by pressing a different button.

**Figure 2.**
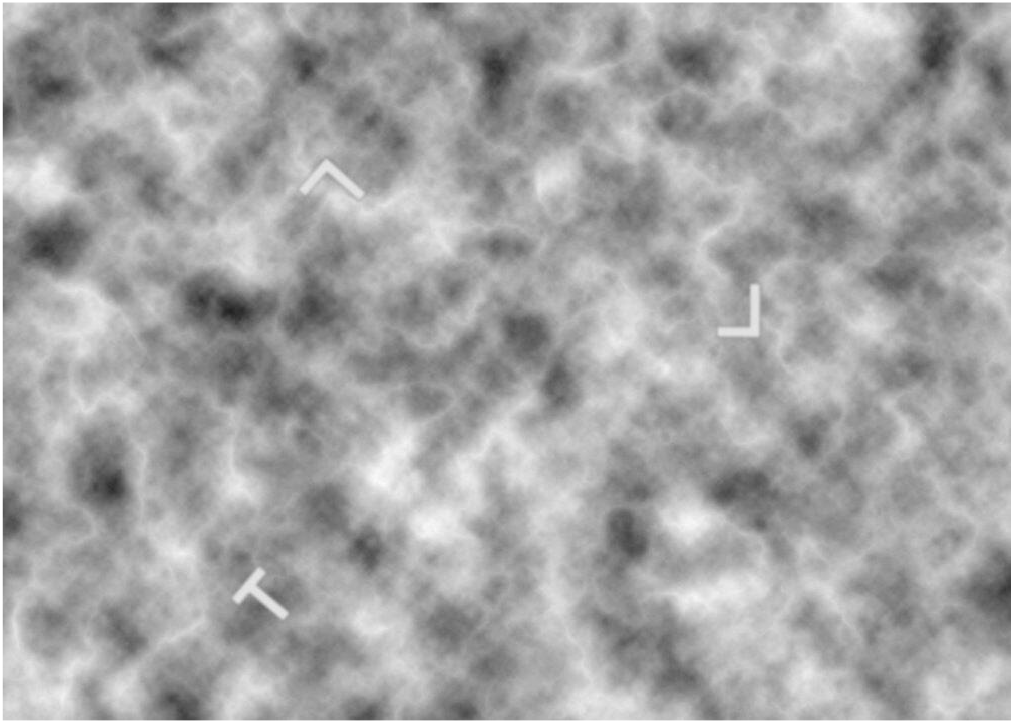
Sample T/L Image. A target ‘T’ is seen in the lower right. In the experiment, a an observational frame may or may not have included a “T” target.

Detection accuracy (FP and misses) was assessed, along with response times, and eye-tracking allowed us to determine whether the subject visually fixated on a target. Subjects performed the task within the MRI bore, and their neurocognitive states were assessed by *f*MRI.

Brain regions measured for the DMN components in the *f*MRI analysis included the ventromedial and dorsomedial prefrontal cortex, the posterior cingulate cortex, the posterior inferior parietal lobule, and the lateral temporal cortex. FPN brain regions included the anterior prefrontal cortex, dorsolateral prefrontal cortex, anterior cingulate, anterior inferior parietal lobule, and anterior insular cortex. There was some overlap of subjects between these three experiments, with three subjects having participated in at least 2 of the experiments and one subject having participated in all three.

A third experiment used *f*NIRs alone (NIRx Scout, NirX Medical Technologies, LLC, Berlin) and took place in the actual clinical setting. This was an observational study using 9 radiologists (4 men and 5 women) to verify the feasibility of using portable *f*NIRs technology to assess radiologists’ neurocognitive states as they performed their usual interpretive work tasks. We determined what fraction of the radiologists’ time was in the high-risk EPS target-state as they worked, using the same *f*NIRs criteria as in the CAT task experiment. The prevalence of EPS was correlated with subject age. As before, the presence of the EPS was inferred when there was simultaneous activation of the ventromedial PFC (a component of the DMN) and the dorsolateral PFC (a component of the FPN). Since it is known that all of the several brain areas that comprise each network demonstrate tightly-coupled, coordinated patterns of activation, such that it would be unusual for some components of either the DMN or FPN to be active when others are not, our rationale was that since the EPS is task-independent, we should observe an overall prevalence of the EPS state in the clinical setting that is similar to *f*MRI and *f*NIRs our other two experiments.

### Statistical Analysis

All statistical modeling for our data analysis was accomplished using SAS Software 9.4 with the GLIMMIX procedure. For our first two experiments, generalized linear modeling (GLM) was used to model state frequency and duration assuming a Poisson and binomial distribution, respectively. False negative rate by state was examined using generalized linear mixed modeling assuming a binary distribution, where observations were nested within radiologists (radiologist were a random effect). In the analysis of the results of Experiment 3, EPS was modeled by age using GLM assuming a binomial distribution. For additional details on methods and technical specifications see the appended technical supplement.

## Results

### (1) Experiment 1: Continuous Attention Task

In the first experiment using the continuous attention task (CAT), the *f*MRI data from one (male) subject was corrupted and could not be analyzed, leaving 9 fMRI subjects for whom excellent quality *f*MRI and task-level data were available for analysis. Of these, there was one (female) subject for which a malfunction degraded the *f*NIRs data, so that technically excellent quality *f*NIRs data were ultimately obtained for 8 subjects. The average error (miss) rate per subject was 10% (range 2% - 26%). Correct response times ranged from 550ms - 2.0 seconds (mean = 1.2 sec). There were a total of 52 FN and 15 FP errors in our 9 subjects.

Irregular, aperiodic brief episodes of EPS were observed, as illustrated with blue shading in Figure 3, below, which is of a single-subject from Experiment 1. During the CAT visual task, on both *f*MRI and *f*NIRs, cortical activation patterns were observed to shift between periods of activity in the FPN, alternating with periods of DMN activation (generally opposed), punctuated by brief, episodes where subjects were observed to be in the target EPS state, *i*.*e*., where both networks were simultaneously active. The number and duration of spontaneous episodes of EPS detected varied, ranging from 6 – 12 in each 15-minute block, with an average of 8.5 spontaneous EPS events, most lasting on the order of 2-4 seconds. Their temporal occurrence appeared to be spontaneous, aperiodic and irregular, with no apparent relationship to the visual task.

**Figure 3.**
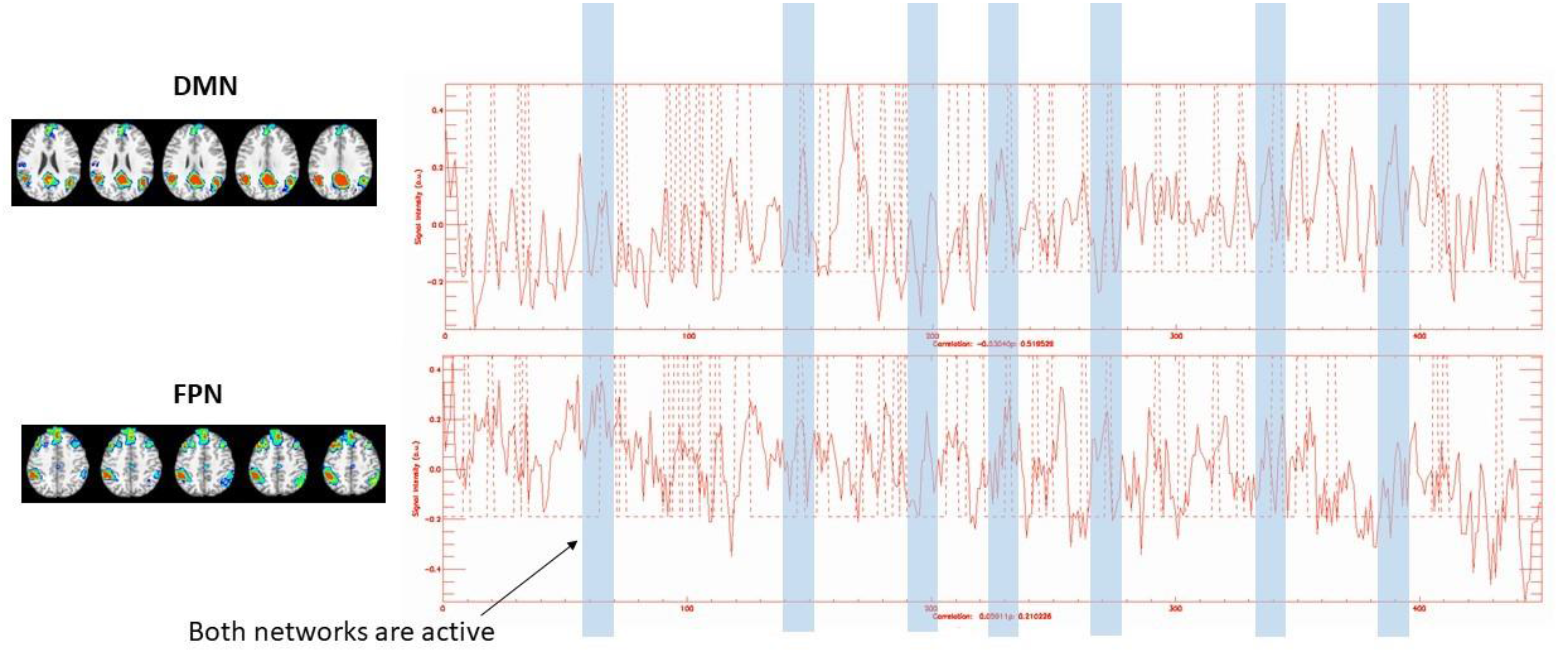
Experiment #1. Representative *f*MRI data from one subject radiologist showing the relative amplitude time course of network activation as a function of time during the 450-second CAT task for DMN (upper) and FPN (lower). Y-axis = relative signal intensity amplitude of network activation by *f*MRI (BOLD) and X-axis = time in seconds (0-450). These two networks were observed to be generally opposed, as expected, however several brief episodes of co-activation of both networks were observed (blue shaded regions). These episodes correspond to the EPS. For this subject, 7 such episodes were detected during the session.

Subjects were found to be in the EPS approximately 20 – 25% of the session time. As predicted, the EPS state was found to correlate strongly with FN errors but not FP errors. Although FN errors were relatively few, and episodes of EPS were relatively brief, a total of 37 FN errors (comprising 71% of the total 52 FN errors) were found to have occurred during EPS episodes by *f*MRI (all 9 subjects) and 29 out of 39 FN errors occurred during EPS by *f*NIRs criteria in the 8 subjects for which *f*NIRs data were collected. This was a highly significant finding, p<0.01 (generalized linear modeling). FN errors were also seen at times of state transition or during periods of DMN activation without FPN activation. These results are tabulated below.

Representative *f*NIRs results from Experiment 1 are shown as Figure 4, below. *f*NIRs was able to spatially discriminate activity between the two adjacent PFC areas (medial vs. lateral PFC) and identify their distinct activation states (figs 3a and 3b) and detect EPS (fig 3c) with a physiologically robust BOLD signal response (fig 3d). There was 73% overall concordance between the *f*MRI results of network activation dynamics and the corresponding *f*NIRs results, including for frequency, timing and duration of EPS episodes, and there was agreement between both modalities in 67% of the detected FN+EPS errors.

**Figure 4.**
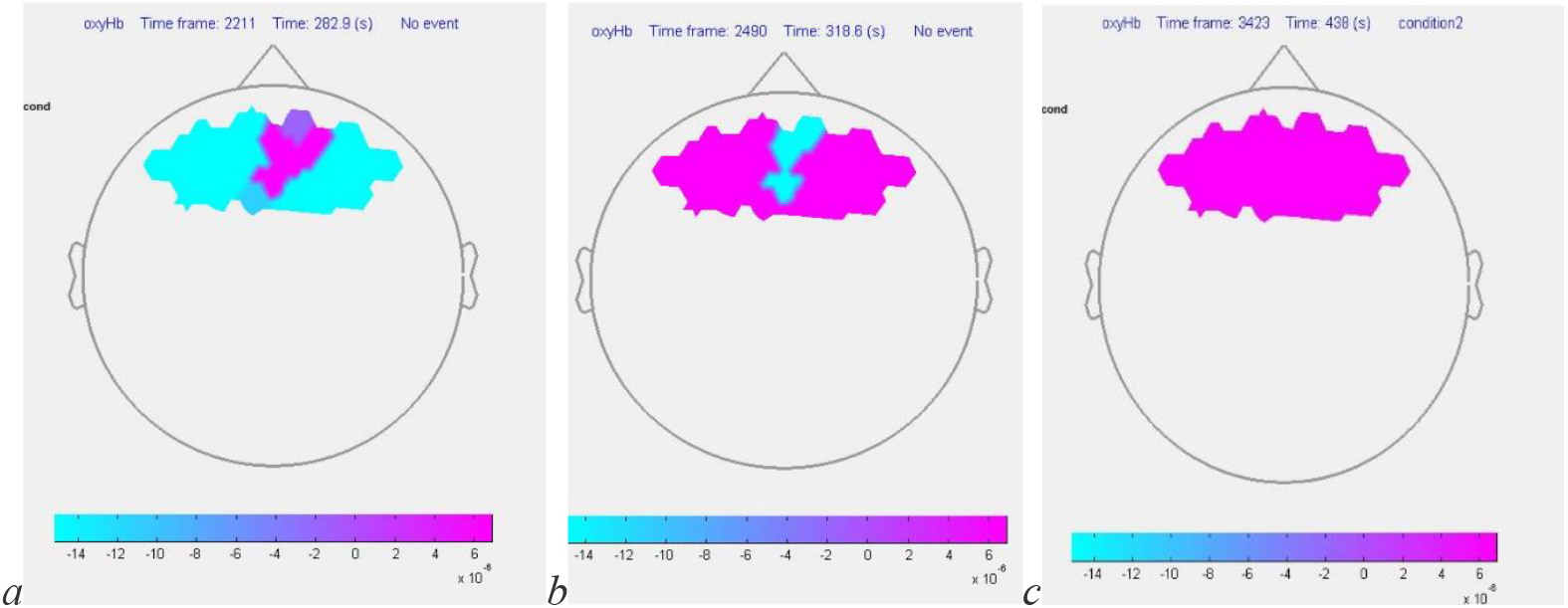
a-c: Experiment 1 *f*NIRs result (above). Representative spatial activation maps from *f*NRIs of BOLD (OxyHb) signal in a single subject, obtained while performing the CAT task while simultaneously undergoing *f*NIRs and *f*MRI. Maps depict signal within the channels corresponding to the medial (a) and dorsolateral (b) PFC activation occurring between tasks, corresponding to fluctuations between DMN and FPN activity as shown on *f*MRI, and (c) in EPS. FN errors (condition 2) were highly correlated with EPS for both *f*MRI and *f*NIRs (p < 0.01).

**Figure 4.**
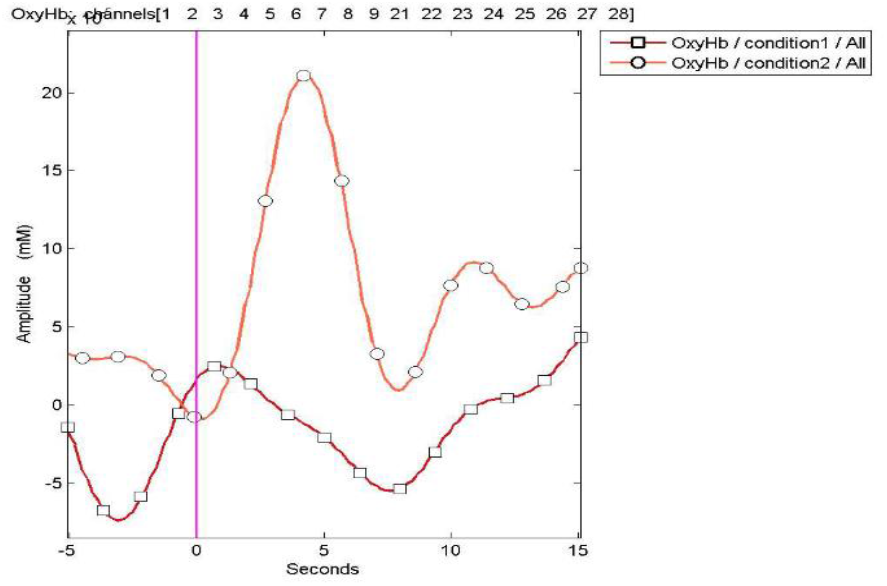
4d (left). Experiment 1. Block-average plot of OxyHb levels from *f*NIRs in one subject corresponding to medial PFC channels of both cerebral hemispheres (DMN) during successful event detections (condition 1) and missed event detections (condition 2), corresponding to an episode of EPS. This plot illustrates the robust discriminatory ability of *f*NIRs to detect physiological activity within selected areas of PFC. Medial PFC areas (DMN) are highly activated during condition 2 (FN errors) compared to condition 1 (TP), as predicted. Dorsilateral PFC (a component of the FPN) was also active at this time. Perceptual errors did not correlate with DMN activation alone.

**Figure 4.**
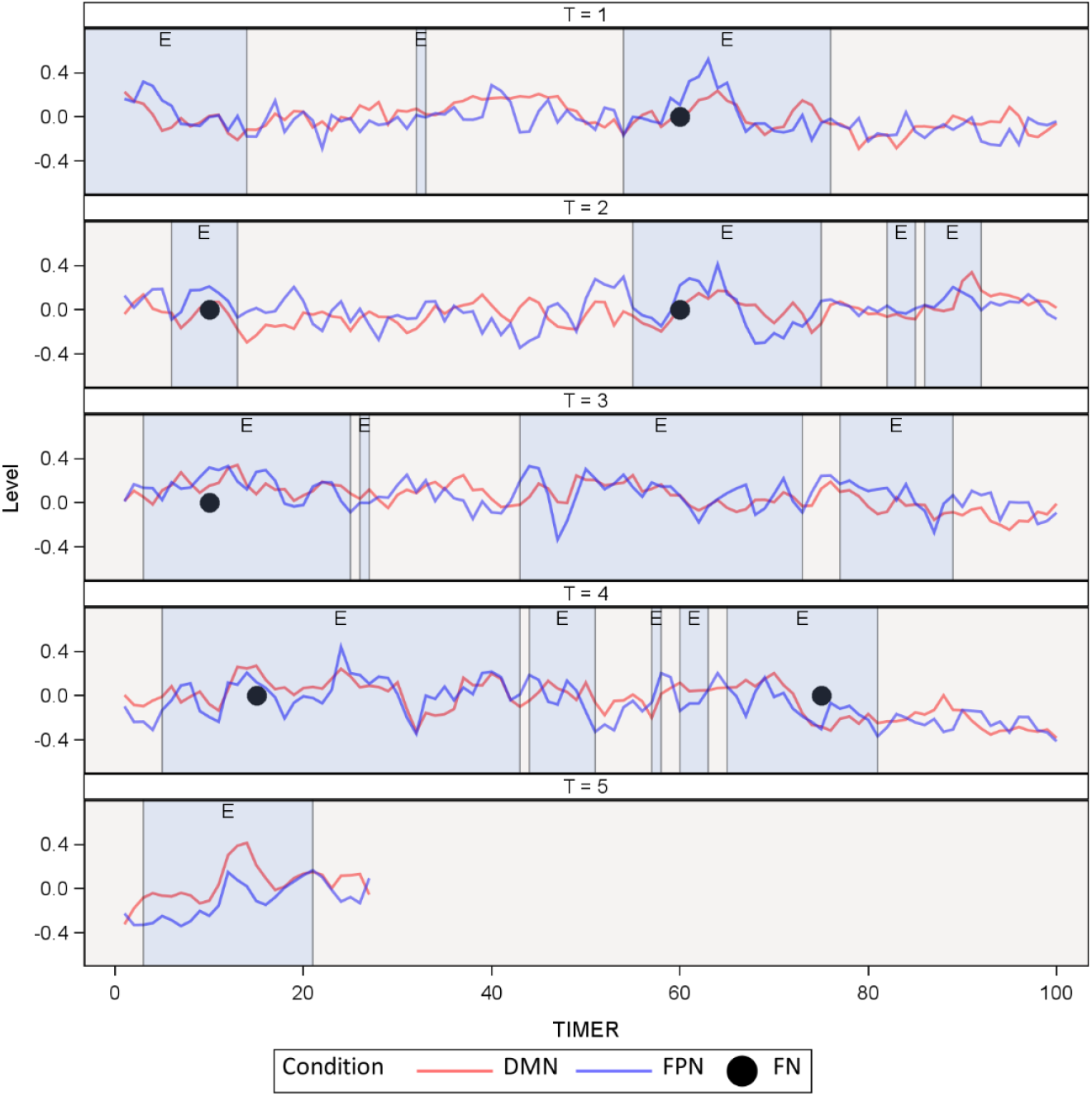
(above) showing results from Experiment 2: Representative *f*MRI data from one radiologist during the T/L search task, showing a 330 second sample within the time course of network activation (*p*<0.01) during a trial session. Curves reflect the levels of activation within the DMN (red) and FPN (blue). When the Pearson Coefficient of correlation between the two networks exceeds the target threshold of 0.4, an EPS event (E, blue-shading) is indicated. As before, activation in each of the two networks was generally anti-correlated, but with several brief and variable episodes of simultaneous co-activation (blue shaded regions) occurring. For this subject, 17 such episodes are shown, of varying durations. Black dots indicate the timing of FN error events, when the subject failed to detect the “T” target. Two of the FN errors were noted to have occurred despite the subject visually fixating on the “T” target. This subject also had two FP errors (not shown) which were unrelated to episodes of EPS. Y-axis = relative amplitude of network activations for DMN and FPN. X-axis = time in seconds (0-400).

### (2) Experiment 2: Visual Search Task, with eye-tracking

In the second experiment, 6 subjects performed a visual search task using eye-tracking with *f*MRI. There were a total of 28 FN and 2FP errors observed, from a total of 360 trials, for an overall error rate of 7.7% across all subjects. Individually, subjects made 4.3 FN errors per session on average (range 2-7). Only one subject experienced any FP errors. Of 28 FN errors, there were 13 where the subjects had visually fixated on the target as shown by eye-tracking. Within a 450-second window containing 60 trials, the average number of EPS episodes per subject was 11 (95% CI, range 9.5-12.7) and the average fraction of time spent in EPS was 28.9% (95% CI, range 24.1% - 34.24%), or approximately 4.3 minutes. In considering the relative risk of FN errors comparing the EPS to non-EPS neurocognitive states, the average FN rate with 60 trials was 10.75% (95% CI, range 5.06 - 21.42) when not in EPS and 30.33% (95% CI, range 19.82 - 43.40) when in EPS, an odds increase of 3.61 (95% CI, range 1.39 - 9.35), with *p*=.0087. If the FN rates are normalized for the relative durations of EPS and non-EPS states, the odds ratio falls slightly, to 2.84. This shows that EPS is a substantial risk factor for FN errors for this task (approximately threefold increased error risk). There was no evidence of any difference between FN errors whether or not eye-tracking showed fixation on the “T” target. There were only two FP errors, however, which both occurred in the same subject. These were unrelated to EPS.

**Table 1:** EPS and FN Errors.

| Experiment | FN in EPS | FN not in EPS | % FN in EPS | n | p-value |
| --- | --- | --- | --- | --- | --- |
| Experiment 1 - fMRI | 37 | 15 | 71% | 9 | $P < 0.01$ |
| Experiment 1 - fNIRs | 29 | 10 | 74% | 8 | $P < 0.01$ |
| Experiment 2 - fMRI | 21 | 8 | 72% | 6 | $P < 0.01$ |

### (3) Experiment 3: Observational study of radiologists in the actual clinical work setting

For the 9 radiologists who were observed using *f*NIRs in the actual clinical setting while engaged in their normal interpretive work activities, subjects were found to be in the EPS on-average 23.5% (range 17-33%, with SD = 5.38) of their working time.

**Table 2:** Fraction of Time in EPS.

| Experiment | Range | Mean | S.D. |
| --- | --- | --- | --- |
| Experiment 1 | 24-35% | 30.4% | 3.8% |
| Experiment 2 | 24-34% | 28.9% | 4.8% |
| Experiment 3 | 17-33% | 23.5% | 5.4% |

These EPS rates are in-line with the existing literature, in which subjects had been found to be in the EPS 20-30% of the time during demanding mental tasks (24) and with the results of our two prior experiments with search and non-search visual tasks using *f*MRI as well as *f*NIRs. There was also a significant anti-correlation of the EPS fraction by subject age (p<0.001), as shown in Figure 5. This finding also aligns with past data (25, 26), suggesting that older adults are off-task less frequently than younger adults. No attempt was made during this experiment to determine whether any interpretive errors were made by any of the subjects during the period of observation.

**Figure 5.**
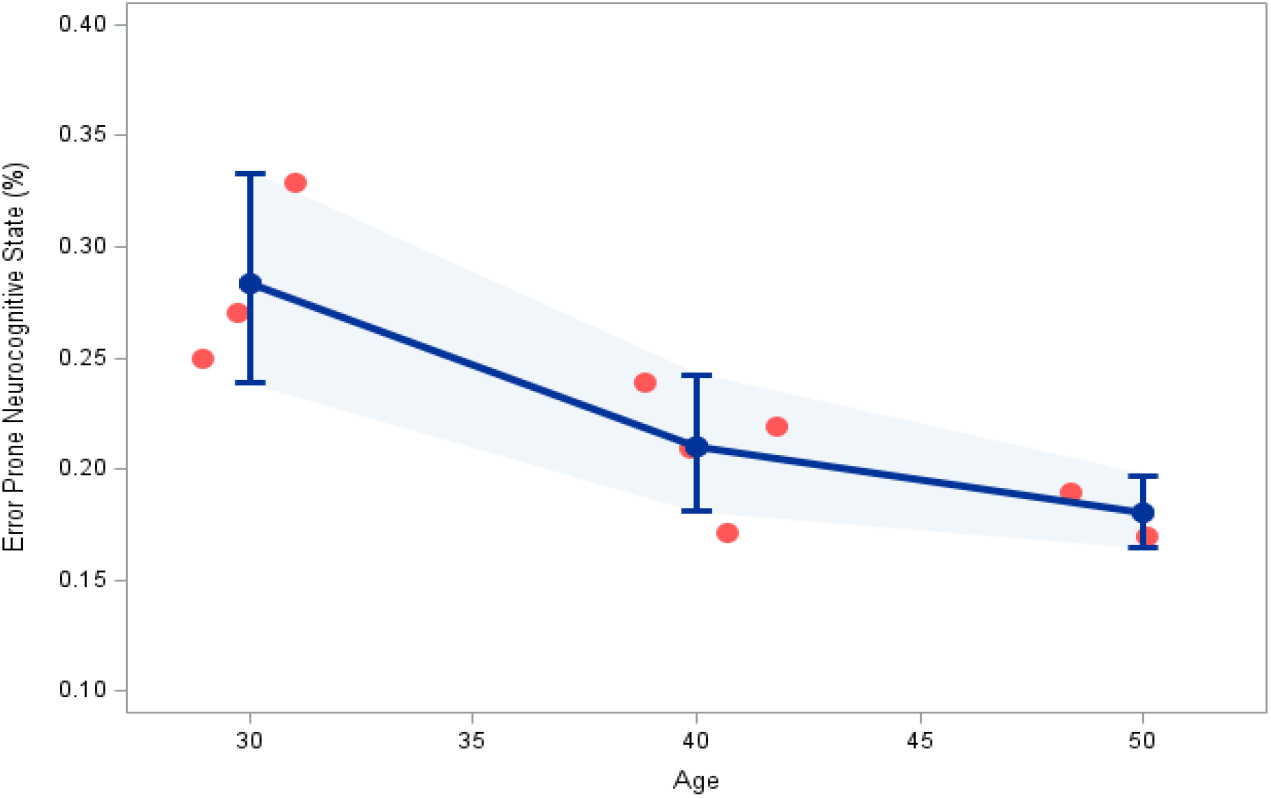
(above): Prevalence of EPS by age, as determined from *f*NIRs of 9 subjects during their normal tasks, performed in the actual clinical setting, showing anti-correlation of EPS prevalence with age (p<0.001). Error bars = 95% CI.

## Discussion

While subject numbers are low in this pilot study, these results support the hypothesis that an error-prone state (EPS) arising from episodic co-activation of the nominally-opposed DMN and FPN networks—which appears to occur spontaneously on a stochastic basis and is non-volitional—is a marker of enhanced error susceptibility for a majority of the observed perceptual errors in our radiologist subjects. FN (miss) errors were found to be significantly more likely in the presence of this neurocognitive EPS than at other times, with an approximately threefold increase in the risk of error at these times. The relationship between the observed EPS and false-negative errors was observed in both a search task and a continuous-attention challenge task. Our results also support the conclusion that the EPS can be detected unobtrusively in the clinical setting.

Our preliminary data suggest that the EPS state represents an involuntary neurocognitive human factor that may be a major contributing mechanism for radiologists’ perceptual errors in practice, which also appear to occur randomly and without the awareness of any attentional lapse or departure from usual image evaluation on the part of the operator. Since some errors are observed in the absence of EPS, it cannot be the sole mechanism of “look-but-fail-to-see” errors, however our data suggest that intervention strategies based on EPS may be able to reduce the prevalence of such errors in practice.

The prevalence of EPS in our relatively small sample is also in keeping with the observed prevalence of perceptual errors in actual radiological practice, which has remained essentially unchanged since first measured in the late 1940s (6). This mechanism of error would also account for the observed higher error rates occurring when the case mix is artificially “enriched” with a larger fraction of positive cases than are usually seen in routine practice.

The EPS episodes we observed, using two independent measures of BOLD signal, were found to be variable in duration and frequency, but occurred with similar prevalence across all tasks studied, including when radiologists performed their usual image interpretation tasks in their actual clinical practice work settings. We found that the unobtrusive, portable technique of *f*NIRs correlated with the “gold standard” of *f*MRI to a high degree, with a 73% concordance in frequency, timing and duration of EPS events and concordance on 67% of FN errors observed during EPS, using the analytical criteria described. In this way, our study also has demonstrated the feasibility of using the unobtrusive method of *f*NIRs to reliably and reproducibly (if imperfectly) detect the EPS in the actual clinical work setting, providing a pathway for intervention to reduce the risk of perceptual errors in practice.

Our finding of a high level of concordance between *f*MRI and *f*NIRs is consistent with very recently published results using concurrent whole-brain *f*MRI and *f*NIRs using similar methods in a different experimental context (27). We would not have expected a perfect correlation between the two modalities, since *f*NIRs assesses only two brain areas compared to several for *f*MRI, however the high level of correlation that we observed between the two modalities should be more than sufficient to allow the development and testing of intervention strategies based on *f*NIRs in future work. Further, there is some evidence that certain subsystems within the DMN, including the medial PFC, may have a greater contribution to attentional lapses and thus the risk of perceptual error in radiology than others, which, if correct, would make the approach we envision potentially even more effective (28, 29). Distinct interactions between various individual components of the FPN and DMN, especially the PFC components, have also recently been implicated in mediating people’s level of conscious awareness (30, 31).

We have focused on FN “miss” errors, as these account for the overwhelming majority of radiologists’ errors in practice (2,3,4), and indeed we have again found FP errors to be quite rare and, as expected, unrelated to the neurocognitive mechanisms that we are studying. A perhaps surprising result was noted with the addition of eye-tracking to our visual search task: it would appear that there is no difference, from the standpoint of the EPS, between FN errors where the subject foveated the target and when they did not, although the number of observations was not large. It is nonetheless an intriguing result that we did not detect a difference between “not looking” and “looking without perceiving,” which will need further validation with larger sample sizes in subsequent research. It is possible that when not in EPS, FN errors could be related to a subject simply not fixating on the target, whereas in EPS the FN error persists despite fixation.

Further work will also be needed to determine whether interventions based on detection of the EPS from real-time monitoring of neurocognitive states, perhaps aided by an even less obtrusive monitoring system with data-streams assessed in real-time using a deep-learning algorithm, will be effective to reduce the risk of radiologists’ perceptual errors in practice. Biofeedback could be used in a manner similar to that used by Mills, *et. al*. (32), possibly with the addition of unobtrusive pupillometry, as was done by Groot, *et al*. (33). It may ultimately be possible to develop a wearable system that incorporates a multimodal real-time assessment of high error risk-status of observers in real-time, including a combination of AI integration of eye movements (as was done by Mills, *et al*.) or pupillometry (as was done by Groot, *et al*.) and BOLD spectroscopy with *f*NIRs discriminating activity in the prefrontal cortex (as we have done in this work). By combining these modalities, we may reasonably expect to increase both the sensitivity and specificity for detection of the high-risk neurocognitive state, in order to allow rapid biofeedback to be presented to the user. Such user feedback may, in turn, allow radiologists to make behavioral changes in order to reduce the risk of perceptual error resulting from this neuro-cognitive mechanism. Further work will be needed to determine whether a biofeedback approach as envisioned will meaningfully reduce perceptual error, and whether suitably unobtrusive wearable devices can be fabricated to facilitate this purpose. Intervention strategies could also potentially utilize machine-learning algorithms trained to recognize the EPS signal from a wearable device in real-time, and then simply re-present the portions of any imaging study that the operator was viewing at the time of an EPS event, and thereby avoid the risk of creating “alarm fatigue” on the part of the user.

The key limitation of this preliminary work is the small sample size and relative paucity of error observations, which must be addressed in future studies. Other limitations include our recruitment of subjects who are all highly experienced radiologists, as results from individuals with less training and experience (*e*.*g*., residents or fellows) or even laypersons, may differ. The predominantly male, and predominately right-handed subject pool in our preliminary studies is also a limitation, as there may be gender and handedness differences in EPS. Finally, although we did not strictly control for time of day and thus have not addressed the potentially confounding effect of fatigue, which is known to affect radiologists to differing degrees across the lifecycle (34) our subject runs were confined to the late morning and early-to-mid afternoon (see technical supplement) in an attempt to minimize this potential confounding variable. Future work is planned to more systematically evaluate the effect of fatigue on the prevalence of EPS and resultant FN error.

In conclusion, further research is also needed to better establish the relationship between perceptual error and brain network states in larger samples of subjects, including laypersons and radiologists, and with a broader range of tasks with varying levels of task difficulty. It is encouraging that our results are both highly significant and align well with previously published findings in laypersons performing a variety of visually-based tasks, suggesting that we are observing a normal feature of human higher mental function. As noted above, while the anti-correlation of EPS prevalence with age that we observed has also been reported previously in laypersons, in our sample this may also relate to the radiologists’ intensive training and long experience in performing concentrated visual search and discrimination tasks, *i*.*e*., a “practice effect” producing a lesser degree of mental deviation from the task with increasing years of experience. This hypothesis requires further experimental exploration.

## Supporting information

see the appended technical supplement.

## Abbreviations

DMN: Default Mode Network
FPN: Frontoparietal Network
EPS: The “Error-Prone State” observed with simultaneous co-activation of the DMN and FPN, which are generally opposed.
fMRI: functional MRI
fNIRs: fnctional near-infrared spectroscopy, an inobtrusive functional brain imaging modality
FN: false-negative errors
FP: false-positive errors

## Acknowledgement

The authors gratefully acknowledge the loan of the *f*NIRs equipment, data-collection and analysis software, and technical support provided by the NIRx Corporation (Berlin, Germany) which made this work possible. The helpful efforts of Mr. Jeffrey Vesek, NMR Core Laboratory Director, and of Dr. Christopher S. Freet, Dept. of Psychiatry and Behavioral Health, Penn State College of Medicine, is also gratefully acknowledged. The authors are also deeply grateful to Dr. Jeremy M. Wolfe for many helpful discussions.

## References

1. Itri, J.N., Tappouni, R.R., McEachern, R.O., Pesch, A.F. and Patel, S.H. “Fundamentals of Diagnostic Error in Imaging” RadioGraphics 2018; 38:1845–1865

2. Bruno, M.A., Walker, E.A. and Abuhudeh, H.H. “Understanding and Confronting our Mistakes: The Epidemiology of Error in Radiology and Strategies for Error Reduction.” RadioGraphics 2015; 35:1668–1676

3. Waite, S., Scott, J., Gale, B., Fuchs, T., Kolla, S., Reede D. “Interpretative error in radiology.” Am J Roentgenol (AJR) 2017; 208, 739–749

4. Kim, Y. and Mansfield, L.T. “Fool me twice: delayed diagnoses in radiology with emphasis on perpetuated errors” Am J Roentgenol (AJR) 2014; 202(3):465–470

5. Garland, L.H. “On the Scientific Evaluation of Diagnostic procedures.” Radiology 1949; 52:309–328

6. Berlin, L. “Accuracy of Diagnostic Procedures: Has it improved over the past five decades?” Am J Roentgenol (AJR) 2007; 188(5):1173–1178

7. Berlin, L., “Radiologic errors, past, present and future” Diagnosis (Berl). 2014; 1(1):79–84

8. Krupinski EA. (2015). “Improving patient care through medical image perception research: Policy Insights” Behav Brain Sciences 2, 74–80

9. Schooler, J. W., Smallwood, J., Christoff, K., Handy, T. C., Reichle, E. D., & Sayette, M. A. (2011). “Meta-awareness, perceptual decoupling and the wandering mind.” Trends in cognitive Sciences, 15(7), 319–326.

10. Smallwood, J., Brown, K., Baird, B., Schooler J., 2012; “Cooperation between the default mode network and the frontal-parietal network in the production of an internal train of thought.” Brain Research 1428(2012): 60–70.

11. Christoff K., Gordon A., Smallwood, J., Smith, R., Schooler, J., 2009; “Experience sampling during MRI reveals default network and executive system contributions to mind wandering.” PNAS, 106(21), 8719–8724

12. Raichle, M., Macleod, A., Snyder, A., Powers, W., Gusnard, D., Shulman, G. “A default mode of brain function.” PNAS 2001; 98(2) 676–682

13. Greicius, M.D., Krasnow, B., Reiss, A. and Menon, V. “Functional connectivity in the resting brain: A network analysis of the default mode hypothesis.” PNAS 2003;100(1):253–258.

14. Vincent, J.L., Kahn, I., Snyder, A.Z., Raichle, M.E., and Buckner, R.L. “Evidence for a Frontoparietal Control System Revealed by Intrinsic Functional Connectivity.” J Neurophysiol 2008; Dec; 100(6):3328–3342

15. Spreng, R.N., Sepulcre, J., Turner, G.R., Stevens, W.D., Schacter, D.L. “Intrinsic architecture underlying the relations among the default, dorsal attention, and frontoparietal control networks of the human brain.” Journal of Cognitive Neuroscience 2013; 25:74–86

16. Kam, J. W., & Handy, T. C. (2018). Electrophysiological evidence for attentional decoupling during mind-wandering. The Oxford handbook of spontaneous thought: Mind-wandering, creativity, and dreaming, 249–258.

17. Weissman, D.H., Roberts, K., Visscher, K., and Woldorff, M.G. “The neural basis of momentary lapses in attention.” Nat Neurosci 2006; 9:971–978

18. Smallwood, J., Brown, K., Baird, B., Schooler, J. “Cooperation between the default mode network and the frontal-parietal network in the production of an internal train of thought.” Brain Research 2012; 1428(2012): 60–70

19. Kane, M., Brown, L., McVay, J., Silvia, P., Myin-Germeys, I., Kwapil, T. “For whom the mind wanders, and when: An experience-sampling study of working memory and executive control in daily life.” Psychological Science 2007; 18(7):614–621

20. Mason, M., Norton, M., Van Horn, J., Wegner, D., Grafton, S., Macrae C. “Wandering minds: The default network and stimulus-independent thought.” Science 2007; 315(5810): 393–395

21. Reiche, E.D., Reineberg, A.E., Schooler, J.W. “Eye movements during mindless reading.” Psych Science 2010; 21: 1300–1310

22. Mittner, M., Boekel, W., Tucker, A.M., Turner, B.M.,Heathcote, A., Forstmann, B.U. “When the Brain Takes a Break: A Model-Based Analysis of Mind Wandering.” J Neuroscience 2014; 34(49):16286–16295

23. Baird, B., Smallwood, S., Lutz, A., and Schooler, J.W. “The decoupled mind: Mind-wandering disrupts cortical phase-locking to perceptual events.” J Cognitive Neuroscience 2014; 26:2596–2607

24. Eichle, T., Debener, S., Calhoun, V.D., Specht, K., et al. “Prediction of human errors by maladaptive changes in event-related brain networks” PNAS 2008; 105:6173–6178

25. Maillet, D., Beaty, R. E., Jordano, M. L., Touron, D. R., Adnan, A., Silvia, P. J., … & Kane, M. J. “Age-related differences in mind-wandering in daily life.” Psychology and Aging 2018; 33(4), 643

26. Zavagnin, M., Borella, E., & De Beni, R. “When the mind wanders: Age-related differences between young and older adults.” Acta Psychologica 2014; 145, 54–64.

27. Sanchez-Alonzo, S., Canale, R.R., Nicholson, I.F. and Aslin, R.N. “Simultaneous data collection of fMRI and fNIRs Measurements Using a whole-Head Optode Array and Short-Distance Channels.” J. Vis. Exp. 2023; 200:e65088. doi:10.3791/65088(2023)

28. Kucyi, A. “Just a thought: How mind-wandering is represented in dynamic brain connectivity.” Neuroimage 2018; 180:505–514. doi: 10.1016/j.neuroimage.2017.07.001

29. Andrews-Hanna, J.R., Reidler, J.S., Sepulcre, J., Poulin, R., and Buckner, R.L. “Functional-Anatomic Fractionation of the Brain’s Default Network” Neuron 2010; 65:550–52.

30. Long, J., Xie, Q., Ma, Q., Urbin, M.A., Liu, L., et al. “Distinct Interactions between Fronto-Parietal and Default Mode Networks in Impaired Consciousness.” Scientific Reports 2016; doi:10.1038/srep38866 (Nature.com/scientificreports).

31. Dixon, M.L., De La Vega, A., Mills, C., Andrews-Hanna, J., Spreng, R.N., et al. “Heterogeneity within the frontoparietal control network and its relationship to the default and dorsal attention networks.” PNAS 2018; E-1598-E1607 doi: 10.1073/pnas.1715766115

32. Mills, C., Gregg, J., Bixler, R., & D’Mello, S. K. “Eye-mind reader: An intelligent reading interface that promotes long-term comprehension by detecting and responding to mind wandering.” Human– Computer Interaction 2021; 36(4), 306–332.

33. Groot, J.M., Boayue, N.M., Csifcsak, G, Boekel, W. Huster, R., et al. “Probing the neural signature of mind-wandering with simultaneous fMRI-EEG and pupillometry.” Neuroimage 2021; 224: 117412 doi: 10.1016/j.neuroimage.2020.117412

34. Stec, N., Arje, D., Moody, A.R., Krupinski, E.A., Tyrell, P.N. “A Systematic Review of Fatigue in Radiology: Is It a Problem?” Amer. J. Rotentgenol. (AJR) 2018; 210(4):799–806

