## Supplementary material for "Neurocognitive Mechanism of Radiologists’ Perceptual Errors: Results of Preliminary Studies": see the appended technical supplement.

Technical Supplement to

This supplement will provide additional technical details regarding equipment and methods used.

**Functional MRI (*f*MRI)** was performed on a 3.0T Siemens PrismaFit system with a 64-channel head coil and two-channel transmit capability (Erlangen, Germany). Computational support includes a 32-processor cluster. An Eloquence™ System for functional MR imaging (Invivo Corp, FL, USA) including an LCD visual display, audio system, button-response unit and e-prime software for paradigm creation and visual display, subject management, precise delivery of stimuli and behavioral data analysis (Brain Voyager Package). The system is also equipped with an EyeLink 1000 Plus System (SR Research, Ottawa, ON, Canada). The SR Eyelink Host PC performs real-time eye-tracking at 250, 500, 100, or 2000 samples per second while computing true gaze position on the display viewed by the participant. The system also performs analysis of eye-motion events, such as saccades, blinks and fixations.

*f*MRI acquisition parameters utilized a TR of 8.6 ms, TE of 4.00 ms, with a 7mm slice thickness, with 2 averages and a flip angle of 20 degrees, with a phase resolution of 91% and a bandwidth of 320 Hz/Px. Resting-state acquisition was done for several seconds prior to onset of the visual paradigm.

**Functional Near Infrared Spectroscopy (*f*NIRs)** was performed using a NIRx Scout™ system (NIRx Medical Technologies, LLC, Berlin, Germany), using a fully MRI-compatible array of optodes. This system was provided on-loan by the company. *f*NIRs is a well-established noninvasive and portable means to perform functional brain imaging using the BOLD effect, based on the absorption of light in the near-infrared spectrum, from 700 to 1300 nm wavelength. Biological tissues (scalp, skull, meninges) are relatively transparent at these wavelengths, and so the absorption spectra are largely dependent on the oxygenation and deoxygenation states of hemoglobin and the redox state of cytochrome c oxidase in mitochondria. *f*NIRs can measure concentration changes in oxygenated (oxy-Hb) and deoxygenated (deoxy-Hb) hemoglobin within the human brain with high accuracy over short time-frames, and the technique is commonly used in neuroimaging studies and also in limited clinical applications. The basic principle is similar to *f*MRI, however the spatial resolution is significantly lower and signal can only be derived from more superficial brain areas within the optical path, generally at depths less than 25mm. For both experiments 1 and 3, an optode array optimized for the prefrontal cortex was utilized, with 8 laser light sources and 11 detectors, held to the scalp in standard position by a specialized cap designed for this purpose. **Figure 1** (right) illustrates the arrangement of these elements, which is often referred to as the optode *montage*. Data were acquired using NIRStar™ software and analyzed using NIRSlab™ a Matlab-based application developed for this purpose SUNY Downstate and licensed by

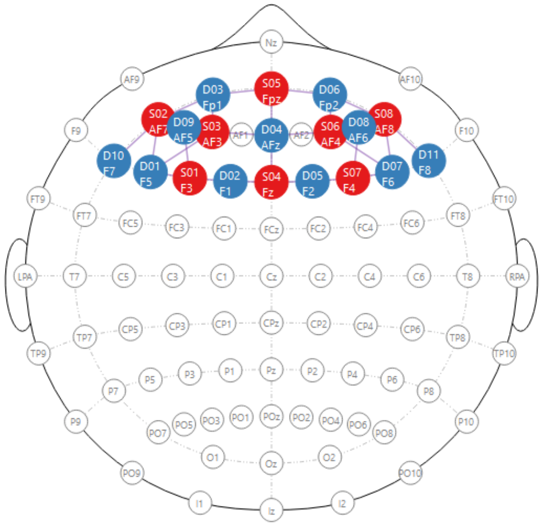


**Figure 1: The 'Optode Montage" showing the physical arrangement of laser light sources (red dots) and detectors (blue dots) used in our fNIRs experiments.**

NIRx (which was provided for our pilot study free of charge). For a full review of *f*NIRs in the context of neuroimaging, see Hoshi, Y., "Hemodynamic signals in *f*NIRs," Chapter 7 in *Progress in Brain Research* 2016; Vol 225, pages 153-179, ISSN 0079-6123, doi: 10.1016/bs.pbr.2016.03.004 (The interested reader will also note that this chapter includes a very comprehensive bibliography of background reading, with 134 references).

**In Experiment 1** we performed *f*MRI and *f*NIRs *simultaneously* as subjects performed the continuous-attention (CAT) task. A signal from the e-prime system launched the visual paradigm and synchronized data acquisition for both the *f*MRI and *f*NIRs systems, functioning independently. The CAT task itself required the subjects to engage for 15 minutes, divided into two 450-second sessions, requiring subjects' continuous attention for 7.5 minutes at a time. All subjects' sessions were scheduled to begin between 11:00 a.m. and 4:00 p.m., to minimize the potentially confounding effects of diurnal / workday fatigue. The subjects were asked to view a grayscale disc on a grayscale background as it slowly meandered through their field of view. At pseudorandom times, and without warning, the disk's luminosity would subtly change. When the subject detected the change, they would signal the detection by a button-push, which was recorded with approximately 10 msec precision, allowing us to measure subject response times. The event is very easy to detect in real-time, but is essentially impossible to identify in retrospect if the subjects' attention had lapsed. The fMRI data were analyzed independently of the *f*NIRs data, using dedicated analytical software developed by one of the investigators (PRK) for this purpose. It quantifies regional brain activity on a relative scale for the pre-identified cortical brain regions which comprise the two targeted networks, the DMN and FPN, in a manner similar to prior publications (see Karunanayaka, P.R., Wilson, D.A., Tobia, M.J., Martinez, B.E., *et al.* "Default Mode Network Activation during Odor-Visual Association" *Human Brain Mapping* 2017; 38:1125-1129 doi 10.1003/hbm.23440). Brain regions chosen to represent the DMN in our analysis included the ventromedial and dorsomedial prefrontal cortex, the posterior cingulate cortex, the posterior inferior parietal lobule, and the lateral temporal cortex. FPN brain regions included the anterior prefrontal cortex, dorsolateral prefrontal cortex, anterior cingulate, anterior inferior parietal lobule, and anterior insular cortex.

Each subject was individually analyzed at the network level, comparing the relative amplitudes of activation of the DMN and FPN networks. Although any degree of positive correlation of the two networks would support our hypothesis, for added rigor, a cutoff value for the Pearson coefficient of correlation between the activity amplitudes of the two networks was set at 0.4. This analysis effectively excluded networks other than the DMN and FPN. For *f*NIRs, BOLD assessment was limited to the prefrontal cortex. This method has less spatial resolution than *f*MRI. We were, however, able to discriminate cortical activation (OxyHb signal) in the *medial PFC*, which is a component of the DMN, from the *dorsolateral PFC*, which is a component of the FPN. When high-amplitude OxyHb signal on the order of 10-20 mM of oxyHb was seen in both PFC cortical regions, it was scored as an EPS event. BOLD signal was measured within these distinct PFC areas during the antecedent 10 sec prior to each visual task event, which is within the temporal resolution of the *f*NIRs method.

In this way, we were able to (1) determine when attentional lapses had occurred (2) determine the underlying brain states at those times to determine whether the EPS (co-activation of both DMN + FPN) was present, on either *f*MRI and/or *f*NIRs, and (3) compare the two. It is known from previously published work that all of the several brain areas that comprise each cortical network demonstrate tightly coupled, coordinated patterns of activation, such that it would be unusual for some to be active while others were not. Based on this, we expected that the *f*NIRs sampling of only one of several components of each network (medial PFC for the DMN and dorsolateral PFC for the FPN) would correlate well with the whole-brain multinuclear analysis done by *f*MRI. One aim of Experiment 1 was to evaluate the level of agreement between the two modalities, given that *f*NIRs is relatively limited. We also determined the fraction of time subjects spent in the EPS during this task, for both the *f*MRI and *f*NIRs.

As an added quality control, in order to verify that the presence of the *f*NIRs system did not degrade the *f*MRI signal, the CAT task was run once with both *f*NIRs and *f*MRI and then again using *f*MRI only, without the *f*NIRs apparatus in place. Signal quality was compared between the two runs to assess whether the *f*NIRs detector system caused any unforeseen artifacts or BOLD signal loss on fMRI. fNIRs data quality metrics (noise and physiological measures) were excellent for both runs.

**Experiment 2** was done using fMRI and eye-tracking, *without* concurrent fNIRs. In this study, the participants were tasked with searching a series of visual images containing multiple distractors for a single target that may or may not have been present. Eye-tracking allowed us to determine whether the targets were foveated by the subjects, and a button-push indicated detection, allowing us to determine whether a detection had successfully occurred, vs. a FN or FP error. fMRI analysis was done using the same software and approach as in Experiment #1. This experiment allowed us to determine the relationship of EPS to both FN and FP errors, and compare results when the target was foveated to when it was not. As before, fMRI acquisition parameters utilized a TR of 8.6 ms, TE of 4.00 ms, with a 7mm slice thickness, with 2 averages and a flip angle of 20 degrees, with a phase resolution of 91% and a bandwidth of 320 Hz/Px. Resting-state acquisition was done for several seconds prior to onset of the visual paradigm. Each subject performed the task twice, in 450-second sessions. Experiment 2 allowed us to evaluate the EPS/Perceptual error hypothesis for the situation of an active visual search by the participant, which differs from the task of continuous attention in Experiment 1. We also determined the fraction of time subjects spent in the EPS during this task.

**Experiment 3** was a trial of *f*NIRs used portably in the actual clinical setting, to determine the fraction of time subject radiologists would be observed to be in the EPS while carrying out their usual interpretive work in the actual clinical setting, with an approximately 15-minute sample, for comparison to Experiments 1 and 2. The purpose of this limited trial was to serve as proof-of-concept that the portable, unobtrusive technique of *f*NIRs could be used in the actual clinical setting to detect EPS for the purpose of real-time intervention, and that the frequency of EPS in this setting would be similar to our other experiments. We also correlated the prevalence of EPS by subject age.

There was some overlap of subjects between these three experiments, with three subjects having participated in at least 2 of the experiments and one subject having participated in all three.

### *Statistical Analysis*

All statistical modeling was accomplished using SAS Software 9.4 with the GLIMMIX procedure. For our first two experiments, generalized linear modeling (GLM) was used to model state frequency and duration assuming a Poisson and binomial distribution, respectively. False negative rate by state was examined using generalized linear mixed modeling assuming a binary distribution, where observations were nested within radiologists (radiologist were a random effect). For Experiment 3, EPS was modeled by age using GLM assuming a binomial distribution.
